# Remembrance of Inferences Past

**DOI:** 10.1101/231837

**Authors:** Ishita Dasgupta, Eric Schulz, Noah D. Goodman, Samuel J. Gershman

## Abstract

Bayesian models of cognition assume that people compute probability distributions over hypotheses. However, the required computations are frequently intractable or prohibitively expensive. Since people often encounter many closely related distributions, selective reuse of computations (amortized inference) is a computationally efficient use of the brain’s limited resources. We present three experiments that provide evidence for amortization in human probabilistic reasoning. When sequentially answering two related queries about natural scenes, participants’ responses to the second query systematically depend on the structure of the first query. This influence is sensitive to the content of the queries, only appearing when the queries are related. Using a cognitive load manipulation, we find evidence that people amortize summary statistics of previous inferences, rather than storing the entire distribution. These findings support the view that the brain trades off accuracy and computational cost, to make efficient use of its limited cognitive resources to approximate probabilistic inference.

## Remembrance of Inferences Past

*“Cognition is recognition.”*

Hofstadter (1995)

## Introduction

Many theories of probabilistic reasoning assume that human brains are equipped with a general-purpose inference engine that can be used to answer arbitrary queries for a wide variety of probabilistic models (Griffiths, Vul, & Sanborn, 2012; Oaksford & Chater, 2007). For example, given a joint distribution over objects in a scene, the inference engine can be queried with arbitrary conditional distributions, such as:

- What is the probability of a microwave given that I’ve observed a sink?
- What is the probability of a toaster given that I’ve observed a sink and a microwave?
- What is the probability of a toaster and a microwave given that I’ve observed a sink?

The nature of the inference engine that answers such queries is still an open research question, though many theories posit some form of approximate inference using Monte Carlo sampling (e.g., Dasgupta, Schulz, & Gershman, 2017; Denison, Bonawitz, Gopnik, & Griffiths, 2013; Gershman, Vul, & Tenenbaum, 2012; Sanborn & Chater, 2016; Thaker, Tenenbaum, & Gershman, 2017; Ullman, Goodman, & Tenenbaum, 2012; Vul, Goodman, Griffiths, & Tenenbaum, 2014).

The flexibility and power of such a general-purpose inference engine trades off against its computational efficiency: by treating each query distribution independently, an inference engine forgoes the opportunity to reuse computations across queries. Every time a distribution is queried, past computations are ignored and answers are produced anew—the inference engine is memoryless, a property that makes it statistically accurate but inefficient in environments with overlapping queries. Continuing the scene inference example, answering the third query should be easily computable once the first two queries have been computed. Mathematically, the answer can be expressed as:

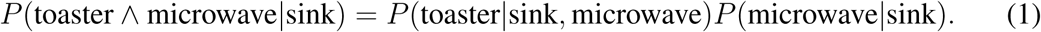

Even though this is a trivial example, standard inference engines do not exploit these kinds of regularities because they are memoryless—they have no access to traces of past computations.

An inference engine may gain efficiency by incurring some amount of bias due to reuse of past computations—a strategy we will refer to as *amortized inference* (Gershman & Goodman, 2014; Stuhlmüller, Taylor, & Goodman, 2013). For example, if the inference engine stores its answers to the “toaster” and “microwave” queries, then it can efficiently compute the answer to the “toaster or microwave” query without rerunning inference from scratch. More generally, the posterior can be approximated as a parametrized function, or *recognition model,* that maps data in a bottom-up fashion to a distribution over hypotheses, with the parameters trained to minimize the divergence between the approximate and true posterior. 1 By sharing the same recognition model across multiple queries, the recognition model can support rapid inference, but is susceptible to “interference” across different queries.

Amortization has a long history in machine learning; the *locus classicus* is the Helmholtz machine (Dayan, Hinton, Neal, & Zemel, 1995; Hinton, Dayan, Frey, & Neal, 1995), which uses samples from the generative model to train a recognition model. More recent extensions and applications of this approach (e.g., Kingma & Welling, 2013; Paige & Wood, 2016; Rezende, Mohamed, & Wierstra, 2014; Ritchie, Thomas, Hanrahan, & Goodman, 2016) have ushered in a new era of scalable Bayesian computation in machine learning. We propose that amortization is also employed by the brain (see Yildirim, Kulkarni, Freiwald, & Tenenbaum, 2015, for a related proposal), flexibly reusing past inferences in order to efficiently answer new but related queries. The key behavioral prediction of amortized inference is that people will show correlations in their judgments across related queries.

We report 3 experiments that test this prediction using a variant of the probabilistic reasoning task previously studied by Dasgupta, Schulz, and Gershman (2017). In this task, participants answer queries about objects in scenes, much like in the examples given above. Crucially, the hypothesis space is combinatorial because participants have to answer questions about sets of objects (e.g., “All objects starting with the letter S”). This renders exact inference intractable: the hypothesis space cannot be efficiently enumerated. In our previous work (Dasgupta, Schulz, & Gershman, 2017), we argued that people approximate inference in this domain using a form of Monte Carlo sampling. Although this algorithm is asymptotically exact, only a small number of samples can be generated due to cognitive limitations, thereby revealing systematic cognitive biases such as anchoring and adjustment, subadditivity, and superadditivity (see also Lieder, Griffiths, Huys, & Goodman, 2017a, 2017b; Vul et al., 2014).

We show that the same algorithm can be generalized to reuse inferential computations in a manner consistent with human behavior. First we describe how amortization might be used by the mind. We consider two crucial questions about how this might be implemented: what parts of previous calculations do people reuse —all previous memories or summaries of the calculations— and when do they choose to reuse their amortized calculations. Next we test these questions empirically. In Experiment 1, we demonstrate that people do use amortization by showing that there is a lingering influence of one query on participants’ answers to a second, related query. In Experiment 2, we explore what is reused, and find that people use summary statistics of their previously generated hypotheses, rather than the hypotheses themselves. Finally, in Experiment 3, we show that people are more likely to reuse previous computations when those computations are most likely to be relevant: when a second cue is similar to a previously evaluated one.

## Hypothesis generation and amortization

Before describing the experiments, we provide an overview of our theoretical framework. First, we describe how Monte Carlo sampling can be used to approximate Bayesian inference, and summarize the psychological evidence for such an approximation. We then introduce amortized inference as a generalization of this framework.

### Monte Carlo sampling

Bayes’ rule stipulates that the posterior distribution is obtained as a normalized product of the likelihood *P(d\h)* and the prior *P(h):*

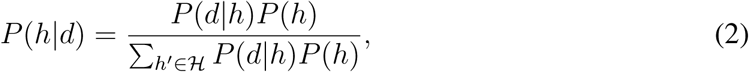

where H is the hypothesis space. Unfortunately, Bayes’ rule is computationally intractable for all but the smallest hypothesis spaces, because the denominator requires summing over all possible hypotheses. This intractability is especially prevalent in combinatorial space, where hypothesis spaces are exponentially large. In the scene inference example, 𝓗 = h_1_ × h_2_ ×…*h_K_* is the product space of latent objects, so if there are *K* latent objects and *M* possible objects, |𝓗| = *M_K_*. If we imagine there are *M =* 1000 kinds of objects, then it only takes *K* = 26 latent objects for the number of hypotheses to exceed the number of atoms in the universe.

Monte Carlo methods approximate probability distributions with samples *θ = {h_1_,…,h_N_*} from the posterior distribution over the hypothesis space. We can understand Monte Carlo methods as producing a recognition model *Q_θ_(h|d)* parametrized by *θ* (see Saeedi, Kulkarni, Mansinghka, & Gershman, 2017, for a systematic treatment). In the idealized case, each hypothesis is sampled from *P(h|d).* The approximation is then given by:

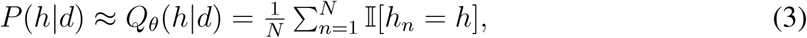

where I[·] = 1 if its argument is true (and 0 otherwise). The accuracy of this approximation improves with *N*, but from a decision-theoretic perspective even small *N* may be serviceable (Gershman, Horvitz, & Tenenbaum, 2015; Lieder et al., 2017a; Vul et al., 2014).

The key challenge in applying Monte Carlo methods is that generally we do not have access to samples from the posterior. Most practical methods are based on sampling from a more convenient distribution, weighting or selecting the samples in a way that preserves the asymptotic correctness of the approximation (MacKay, 2003). We focus on Markov chain Monte Carlo (MCMC) methods, the most widely used class of approximations, which are based on simulating a Markov chain whose stationary distribution is the posterior. In other words, if one samples from the Markov chain for long enough, eventually *h* will be sampled with frequency proportional to its posterior probability.

A number of findings suggest that MCMC is a psychologically plausible inference algorithm. Many implementations use a form of local stochastic search, proposing and then accepting or rejecting hypotheses. For example, the classic Metropolis-Hastings algorithm first samples a new hypothesis 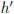 from a proposal distribution 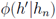 and then accepts this proposal with probability

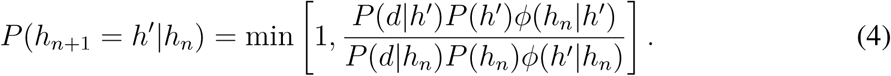

Intuitively, this Markov chain will tend to move from lower to higher probability hypotheses, but will also sometimes “explore” low probability hypotheses. In order to ensure that a relatively high proportion of proposals are accepted, 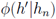 is usually constructed to sample proposals from a local region around *h_n_.* This combination of locality and stochasticity leads to a characteristic pattern of small inferential steps punctuated by occasional leaps, much like the processes of conceptual discovery in childhood (Ullman et al., 2012) and creative insight in adulthood (Suchow, Bourgin, & Griffiths, 2017). Even low-level visual phenomena like perceptual multistability can be described in these terms (Gershman et al., 2012; Moreno-Bote, Knill, & Pouget, 2011).

Another implication of MCMC, under the assumption that a small number of hypotheses are sampled, is that inferences will tend to show anchoring effects (i.e., a systematic bias towards the initial hypotheses in the Markov chain). Lieder and colleagues have shown how this idea can account for a wide variety of anchoring effects observed in human cognition (Lieder, Griffiths, & Goodman, 2012; Lieder et al., 2017b). For example, priming someone with an arbitrary number (e.g., the last 4 digits of their social security number) will bias a subsequent judgment (e.g., about the birth date of Gandhi), because the arbitrary number influences the initialization of the Markov chain.

In previous research (Dasgupta, Schulz, & Gershman, 2017), we have shown that MCMC can account for many other probabilistic reasoning “fallacies,” suggesting that they arise not from a fundamental misunderstanding of probability, but rather from the inevitable need to approximate inference with limited cognitive resources. We explored this idea using the scene inference task introduced in the previous section. The task facing subjects in our experiments was to judge the probability of a particular set of latent objects (the hypothesis, h) in a scene conditional on observing one object (the cue, d). By manipulating the framing of the query, we showed that subjects gave different answers to formally equivalent queries. In particular, by partially unpacking the queried object set (where fully unpacking an object set means to present it explicitly as a union of each of its member objects) into a small set of exemplars and a “catch-all” hypothesis (e.g., “what is the probability that there is a chair, a computer, or any other object beginning with C?”), we found that subjects judged the probability to be higher when the unpacked exemplars were typical (a “subadditivity” effect; cf. Tversky & Koehler, 1994) and lower when the unpacked exemplars were atypical (a “superadditivity” effect; cf. Sloman, Rottenstreich, Wisniewski, Hadjichristidis, & Fox, 2004) compared to when the query is presented without any unpacking.

To provide a concrete example, in the presence of the cue “table,” the typically unpacked query “what is the probability that there is also a chair, a computer, or any other object beginning with C?” generates higher probability estimates relative to the packed query “what is the probability that there is another object beginning with C?”, whereas the atypically unpacked query “what is the probability that there is also a cow, a canoe, or any other object beginning with C?” generates lower probability estimates compared to the packed query.

We were able to account for these effects using MCMC under the assumption that the unpacked exemplars initialize the Markov chain that generates the sample set. Because the initialization of the Markov chain transiently determines its future trajectory, initializing with typical examples causes the chain to tarry in the high probability region of the queried object set, thereby increasing its judged probability (subadditivity). In contrast, initializing with atypical examples causes the chain to get more easily derailed into regions outside the queried object set. This decreases the judged probability of the queried object set (superadditivity). The strength of these effects theoretically diminishes with the number of samples, as the chain approaches its stationary distribution. Accordingly, experimental manipulations that putatively reduce the number of samples, such as response deadlines and cognitive load, moderate this effect (Dasgupta, Schulz, & Gershman, 2017). The experiments reported in this paper build on these findings, using subadditivity and superadditivity in the scene inference paradigm to detect behavioral signatures of amortized inference.

### Amortized inference

As defined in the previous section, Monte Carlo sampling is memoryless, approximating *P(h|d)* without reference to other conditional distributions that have been computed in the past; all the hypothesis samples are specific to a particular query, and thus there can be no cumulative improvement in approximation accuracy across multiple queries. However, a moment’s reflection suggests that people are capable of such improvement. Every time you look out your window, you see a slightly different scene, but it would be wasteful to recompute a posterior over objects from scratch each time; if you did, you would be no faster at recognizing and locating objects the millionth time compared to the first time. Indeed, experimental research has found considerable speed-ups in object recognition and visual search when statistical regularities can be exploited (Oliva & Torralba, 2007).

Amortized inference is a generalization of the standard memoryless framework. We will formulate it in the most general possible terms, and later explore more specific variants.

Figure 1 illustrates the basic idea. In the standard, memoryless framework, an inference engine inverts a generative model *P(d,h)* over hypothesis *h* and data *d* to compute a recognition model *Q_θ_*(h|d) parametrized by θ. For example, Monte Carlo methods use a set of samples to parametrize the recognition model. Importantly, the answer to each query is approximated using a different set of parameters (e.g., independent samples)—*Q_θ1_* (h|d_1_), *Q_θ2_* (h|d_2_), etc. In the amortized framework, parameters are shared across queries. The parameters are selected to accurately approximate not just a single query, but a *distribution* of queries. If cognitive resources are unbounded, then the optimal solution is to parametrize each query separately, thereby recovering the memoryless framework. Under bounded resources, a finite number of parameters must be shared between multiple queries, leading to memory effects: the answer to one query affects the answer to other, similar queries.

**Figure 1.**
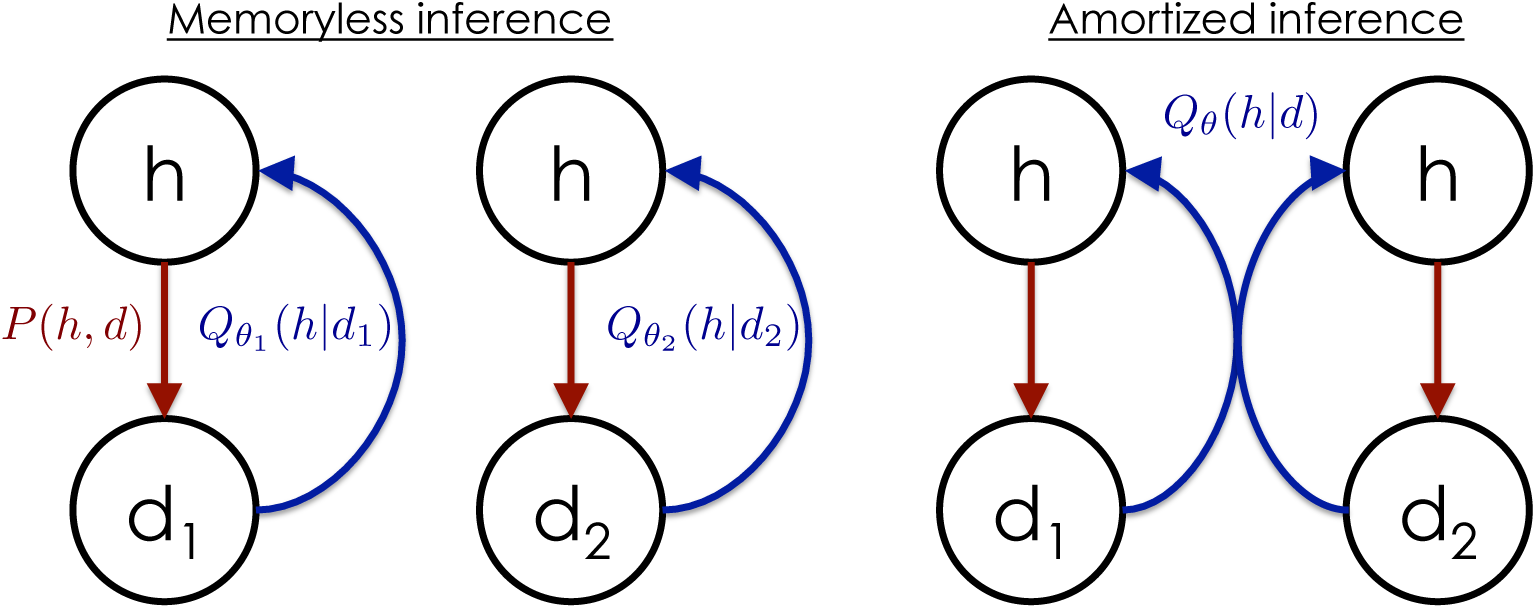
Theory schematic. (Left) Standard, memoryless framework in which a recognition model *Q_θ_(h|d)* approximates the posterior over hypothesis *h* given data *d.* The recognition model is parametrized by *θ* (e.g., a set of samples in the case of Monte Carlo methods). Memoryless inference builds a separate recognition model for each query. (Right) Amortized framework, in which the recognition model shares parameters across queries. After each new query, the recognition model updates the shared parameters. In this way, the model “learns to infer.”

While reuse increases computational efficiency, it can cause errors in two ways. First, if amortization is deployed not only when two queries are identical but also when they are similar, then answers will be biased due to blurring together of the distributions. This is analogous to interference effects in memory. Second, the answer to the first query might itself have been inaccurate or biased, so its reuse will propagate that inaccuracy to the second query’s answer. Our experiments focus on the second type of error. Specifically, we will investigate how the over- or underestimation of unpacked probabilities resulting from approximate inference for one query will continue to influence responses to a second query.

### Two amortization strategies

In our experiments, we ask participants to sequentially answer pairs of queries (denoted *Ql* and *Q2*). In Experiment 2, both queries are conditioned on the same cue object *(d),* but with varying query object sets *(h)*. That is, both questions are querying the same probability distribution over objects, but eliciting the probabilities of different objects in each case. So in theory, all samples taken to answer query 1, can be reused to answer query 2 (they are both samples from the same distribution). This *sample reuse* strategy allows all computations carried out for query 1 to be reused to answer query 2. However, it is expensive, because each sample must be stored in memory. A less memory-intensive solution is to store and reuse summary statistics of the generated samples, rather than the samples themselves. This *summary reuse* strategy offers greater efficiency but less flexibility. Several more sophisticated amortization schemes have been developed in the machine learning literature (e.g., Paige & Wood, 2016; Rezende et al., 2014; Stuhlmüller et al., 2013), but we focus on sample and summary reuse because they make clear experimental predictions, which we elaborate below.

In the context of our experiments, summary reuse is only applicable to problems where the answer to *Q2* can be expressed as the composition of the answer to *Q1* and another (putatively simpler) computation. In Experiment 2, *Q2* queries a hypothesis space that is the union of the hypothesis space queried in *Q1* and a disjoint hypothesis space. For example if *Q1* is “What is the probability that there is an object starting with a C in the scene?”, *Q2* could be “What is the probability that there is an object starting with a C or an R in the scene?”. In this case, samples generated in response to *Q1* are summarized by a single number (“the probability of an object starting with C”), new samples are generated in response to a simpler query (“the probability of an object starting with R”), and these two numbers are then composed (in this case added) to give the final estimate for *Q2* (“the probability of an object starting with C or R”). This is possible because both queries are functions of the same probability distribution over latent objects.

These strategies are simplifications of what the brain is likely doing. Re-using all the samples exactly is unreasonably resource intensive, and re-using only the exact statistic in the few places that the second query can be expressed as a composition of the first query and a simpler computation is unreasonably inflexible. We do not claim that either extreme is plausible, but —to a first approximation— they capture the key ideas motivating our theoretical framework, and more importantly, they make testable predictions which can be used to assess which extreme pulls more weight in controlled experiments.

In particular, sample-based and summary-based amortization strategies make different predictions about how subadditivity and superadditivity change as a function of the sample size (Figure 2, details of these implementations can be found in the Appendix). For sample-based amortization, as the sample size for *Q1* grows, the effect for *Q2* asymptotically *diminishes* and eventually vanishes as the effect of biased initialization in *Q1* washes out. However, initially increasing the sample size for *Q1* also *amplifies* the effects for *Q2* under a sample-based scheme, because this leads to more biased *Q1* samples being available for reuse. The amplification effect dominates up to a sample size of around 230 (estimate for the number of samples taken for inference in this domain, reported in Dasgupta, Schulz, & Gershman, 2017). This effect can be counteracted by increasing the sample size for *Q2*. These are unbiased samples, since *Q2* is always presented as a packed query. More such samples will push the effect down by drowning out the bias with additional unbiased samples.

**Figure 2.**
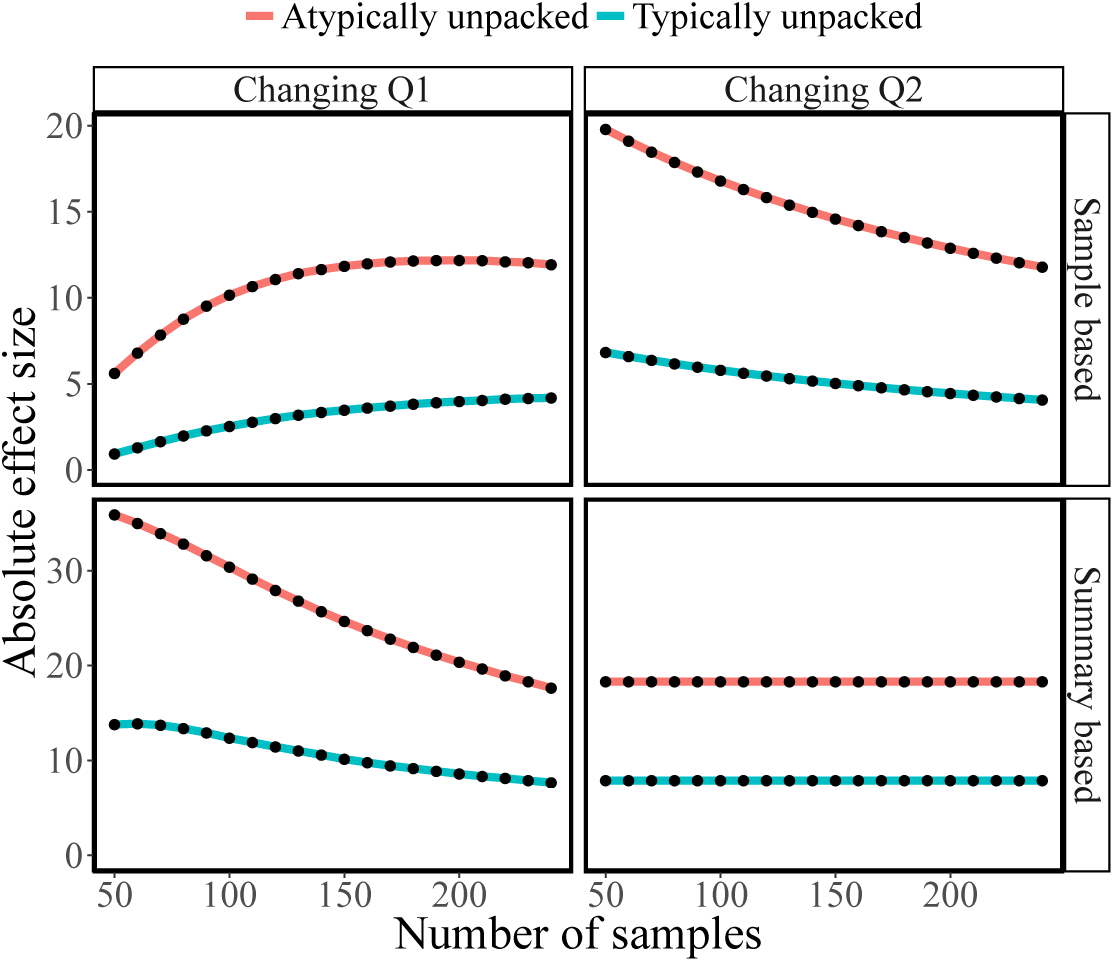
Simulation of subadditivity and superadditivity effects under sample-based (top) and summary-based (bottom) amortization strategies. In all panels, the y-axis represents the unstandardized effect size for *Q2*. Left panels show the effects of changing the sample size for *Q1*; right panels show the effects of changing the sample size for *Q2.* When sample size for one query is changed, sample size for the other query is held fixed at 230 (the sample size estimated by Dasgupta, Schulz, & Gershman, 2017).

Under a summary-based strategy, increasing the sample size for *Q1* will only *diminish* the effects for *Q2*, because the bias from *Q1* is strongest when the chain is close to its starting point. The effect of early, biased samples on the summary statistic disappears with more samples. We see also that changing the number of samples for *Q2* does not influence the effect size because the initialization of the chain for *Q2* is not influenced by the samples or summary statistic from the answer to *Q1*. Under the summary-based strategy, the subadditivity and superadditivity effects for *Q2* derive entirely from the same effects for *Q1*, which themselves are driven by the initialization (see Dasgupta, Schulz, & Gershman, 2017).

We test the different predictions of these strategies by placing people under cognitive load during either *Q1* or *Q2* in Experiment 2, a manipulation that is expected to reduce the number of produced samples (Dasgupta, Schulz, & Gershman, 2017; Thaker et al., 2017). In this way, we can sample different parts of the curves shown in Figure 2.

### Adaptive amortization

Amortization is not always useful. As we have already mentioned, it can introduce systematic bias into probabilistic judgments. This is especially true if samples or summary statistics are transferred between two dissimilar distributions. This raises the question: are human amortization algorithms adaptive? This question is taken up empirically in Experiment 3. Here we discuss some of the theoretical issues.

Truly adaptive amortization requires a method to assess similarities between queries. Imagine as an example the situation in which there is a “chair” in the scene and you have to evaluate the probability of any object starting with a “P”. If afterwards you are told that there is a “book” in another scene, and the task is again to evaluate the probability of any object starting with a “P”, it could be a viable strategy to reuse at least some of the previous computations. However, in order to do so efficiently, you would have to know how similar a chair is to a book, i.e. if they occur with a similar set of other objects on average. One way to quantify this similarity is by assessing the induced posterior over all objects conditioned on either “book” or “chair”, and then comparing the two resulting distributions directly. Cues that are more similar should co-occur with other objects in similar proportions.

To assess the similarity of two distributions over objects induced by two different cues, we will need a formal similarity measure. One frequently used measure of similarity between two probability distribution is the Kullback-Leibler (KL) divergence. For two discrete probability distributions *Q* and *P*, the KL divergence between *P* and *Q* is defined as

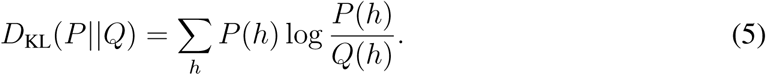

The KL divergence is minimized to 0 when *Q* and *P* are identical. We will use this measure in Experiment 3 to select queries that are either similar or dissimilar, in order to examine whether our participants only exhibit signatures of amortization when the queries are similar.2 Note, however, that the exact calculation of these divergences cannot be part of the algorithmic machinery used by humans to assess similarity, since it presupposes access to the posterior being approximated. Our experiments do not yet provide insight into how humans might achieve tractable adaptive amortization, a problem we leave to future research.

## Experiment 1

In Experiment 1, we seek initial confirmation of our central hypothesis: human inference is not memoryless. To detect these “remembrances of inferences past”, we ask participants to answer pairs of queries sequentially. The first query is manipulated (by packing or unpacking the queried hypothesis) in such a way that subadditive or superadditive probability judgments can be elicited (Dasgupta, Schulz, & Gershman, 2017). Crucially, the second query is always presented in packed form, so any differences across the experimental conditions in answers to the second query can only be attributed to the lingering effects of the first query.

### Participants

84 participants (53 males, mean age=32.61, SD=8.79) were recruited via Amazon’s Mechanical Turk and received $0.50 for their participation plus an additional bonus of $0.10 for every on-time response.

### Procedure

Participants were asked to imagine playing a game in which their friend sees a photo and then mentions one particular object present in the photo (the cued object). The participant is then queried about the probability that another class of objects (e.g., “objects beginning with the letter B”) is also present in the photo.

Each participant completed 6 trials, where the stimuli on every trial corresponded to the rows in Table 1. On each trial, participants first answered *Q1* given the cued object (for example, “I see a lamb in this photo. What is the probability that I also see a window, a wardrobe, a wine rack, or any other object starting with a W?”), using a slider bar to report the conditional probability using values between 0 (not at all likely) to 100 (very likely, see also Figure 3). The *Q1* framing (typical-unpacked, atypical-unpacked or packed) was chosen randomly. Participants then completed the same procedure for *Q2* (immediately after *Q1*), conditional on the same cued object. The framing for *Q2* was always packed and *Q2* was always presented as a conjunction (for example, “What is the probability I see an object starting with a W or F?”), where the order of the letters was determined at random.

**Figure 3.**
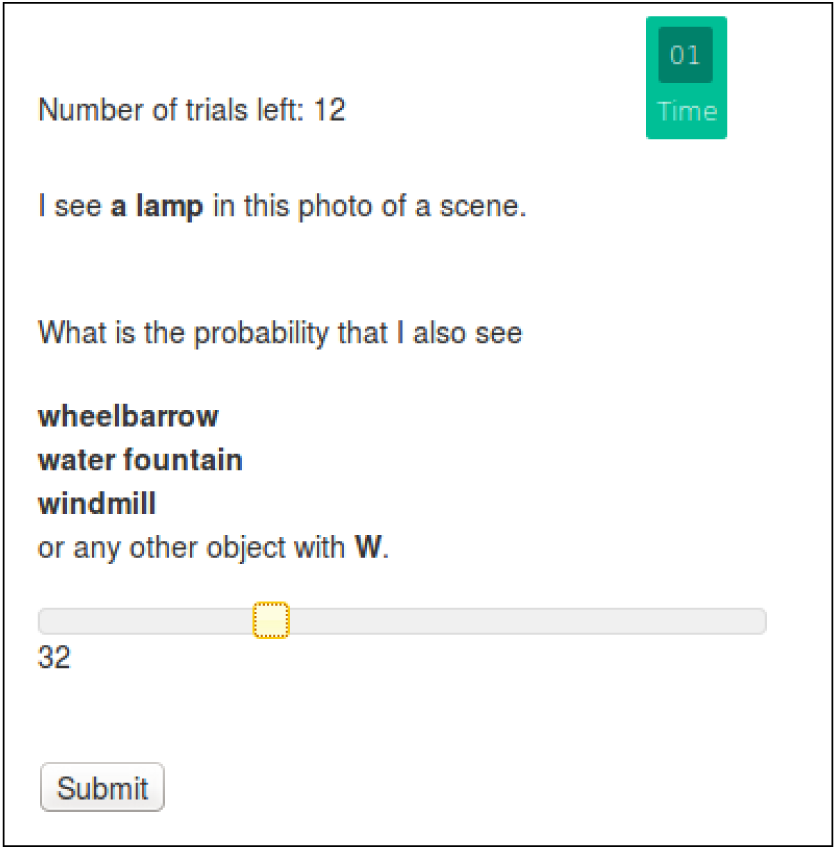
Experimental setup. Participants were asked to estimate the conditional probability using a slider bar within a 20-second time limit.

**Table 1.** Experimental stimuli and queries for Experiment 1.

| Cue | Q1 | Q1-Typical | Q1-Atypical | Q2 |
| --- | --- | --- | --- | --- |
| Table | C | chair, computer, curtain | cannon, cow, canoe | C or R |
| Telephone | D | display case, dresser, desk | drinking fountain, dryer, dome | D or L |
| Rug | B | book, bouquet, bed | bird, buffalo, bicycle | B or S |
| Chair | P | painting, plant, printer | porch, pie, platform | P or A |
| Sink | T | table, towel, toilet | trumpet, toll gate, trunk | T or E |
| Lamp | W | window, wardrobe, wine rack | wheelbarrow, water fountain, windmill | W or F |

### Results

Six participants were excluded from the following analysis, four of which failed to respond on time in more than half of the questions, and two of which entered the same response throughout.

Consistent with our previous studies (Dasgupta, Schulz, & Gershman, 2017), we found both subadditivity and superadditivity effects for *Q1*, depending on the unpacking: probability judgments were higher for unpacked-typical queries than for packed queries (a subadditivity effect; 59.35 vs. 49.67; *t(77)* = 4.03,*p* < 0.001) and lower for unpacked-atypical than for packed queries (a superadditivity effect; 31.42 vs. 49.67; *t(77)* = —6.44,*p* < 0.001).

Next we calculated the difference between each participant’s response to every query and the mean packed response to the same queried object. This difference was then entered as a dependent variable into a linear mixed effects regression with random effects for both participants and queried objects as well as a fixed effect for the condition. The resulting estimates revealed both a significant subadditivity (difference = 12.60 ± 1.25, t(610.49) = 10.083, *p* < 0.0001) and superadditivity (difference = —15.69 ± 1.32, t(615.46) = —11.89, *p* < 0.0001) effect.

Additionally, we found evidence that participants reused calculations from *Q1* for *Q2*: even though all *Q2* queries were presented in the same format (as packed), the estimates for that query differed depending on how *Q1* was presented. In particular, estimates for *Q2* were lower when *Q1* was unpacked to atypical exemplars (46.38 vs 56.83; t(77) = 5.08, *p* < 0.001), demonstrating a superadditivity effect that carried over from one query to the next. We did not find an analogous carry-over effect for subadditivity (58.47 vs. 56.83; t(77) = 0.72, *p* = 0.4), possibly due to the subadditivity effect “washing out” more quickly (i.e. with fewer samples) than superadditivity, as has been observed in this domain before (see Dasgupta, Schulz, & Gershman, 2017).

We calculated the difference between each participant’s response for every *Q2* and the mean response for the same object averaged over all responses to *Q2* conditional on *Q1* being packed. The resulting difference was again entered as the dependent variable into a linear mixed effects regression with both participants and cued object as random effects as well as condition as a fixed effect. The resulting estimates showed both a significant subadditivity (difference = 4.39 ± 1.14, £(606.40) = 3.83, *p* < 0.001) and superadditivity (difference = -7.86 ± 1.21, £(610.41) = -6.50, *p* < 0.0001) effect.

We calculated each participant’s mean response to all packed hypotheses for *Q2* over all trials as a baseline measure and then assessed the difference between each condition’s mean response and this mean packed response. This resulted in a measure of an average effect size for the *Q2* responses (how much each participant under- or overestimates different hypotheses as compared to an average packed hypothesis). Results of this calculation are shown in Figure 4.

**Figure 4.**
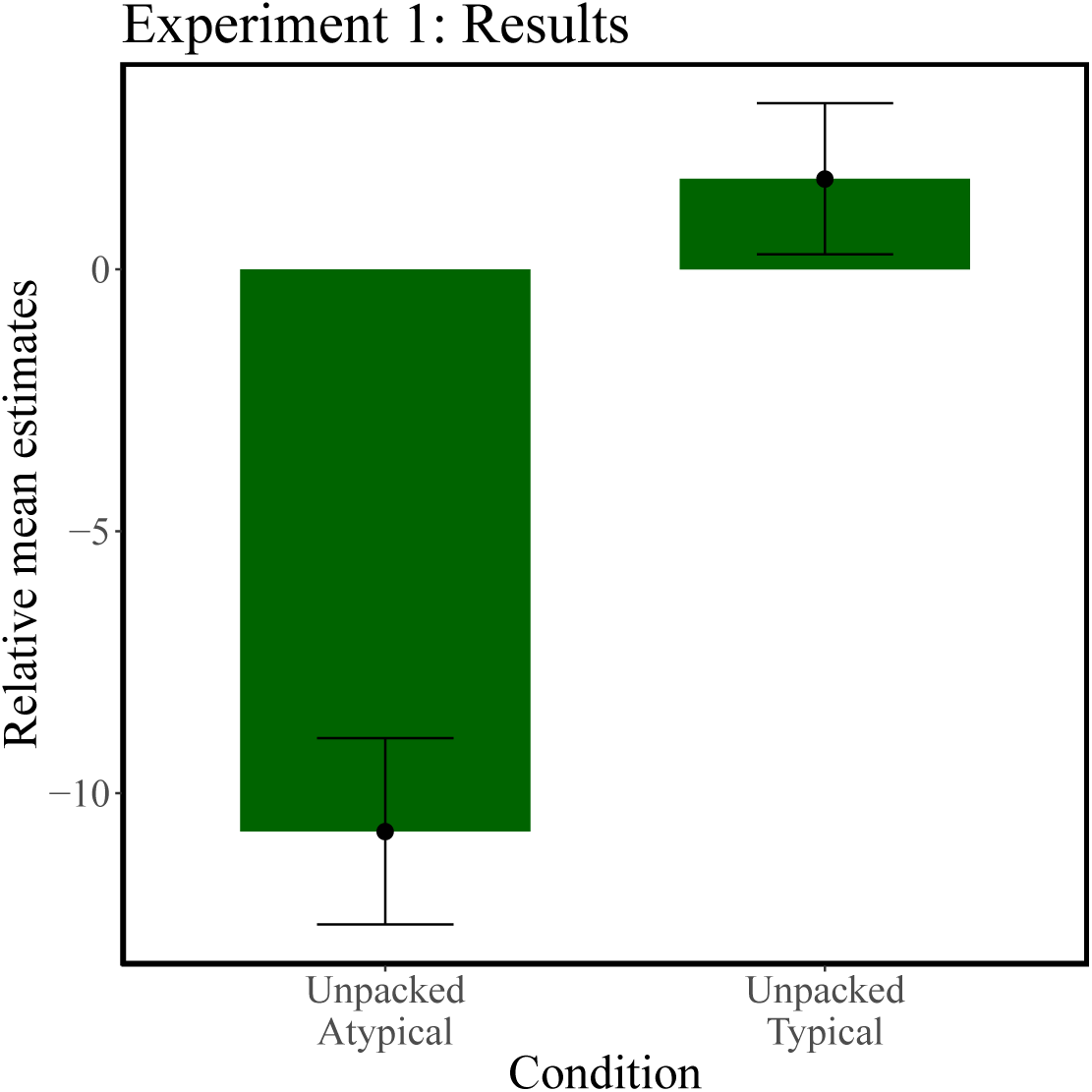
Experiment 1: Differences between *Q2* responses for each condition and an average packed baseline. A negative relative mean estimate indicates a superadditivity and a positive relative mean estimate a subadditivity effect. Error bars represent the standard error of the mean.

The superadditivity effect was significantly greater than 0 (£(77) = 5.07, *p* < 0.001). However, the subadditivity effect did not differ significantly from 0 (£(77) = —0.42, *p* > 0.6; see also Dasgupta, Schulz, & Gershman, 2017).

Next, we explored whether responses to *Q1* predicted trial-by-trial variation in responses to *Q2*. Figure 5 shows the difference between participants’ estimates for *Q1* and the true underlying probability of the query (as derived by letting our MCMC model run until convergence) plotted against the same difference for *Q2*. If participants do indeed reuse computations, then how much their estimates deviate from the underlying truth for *Q1* should be predictive for the deviance of their estimates for *Q2*.

**Figure 5.**
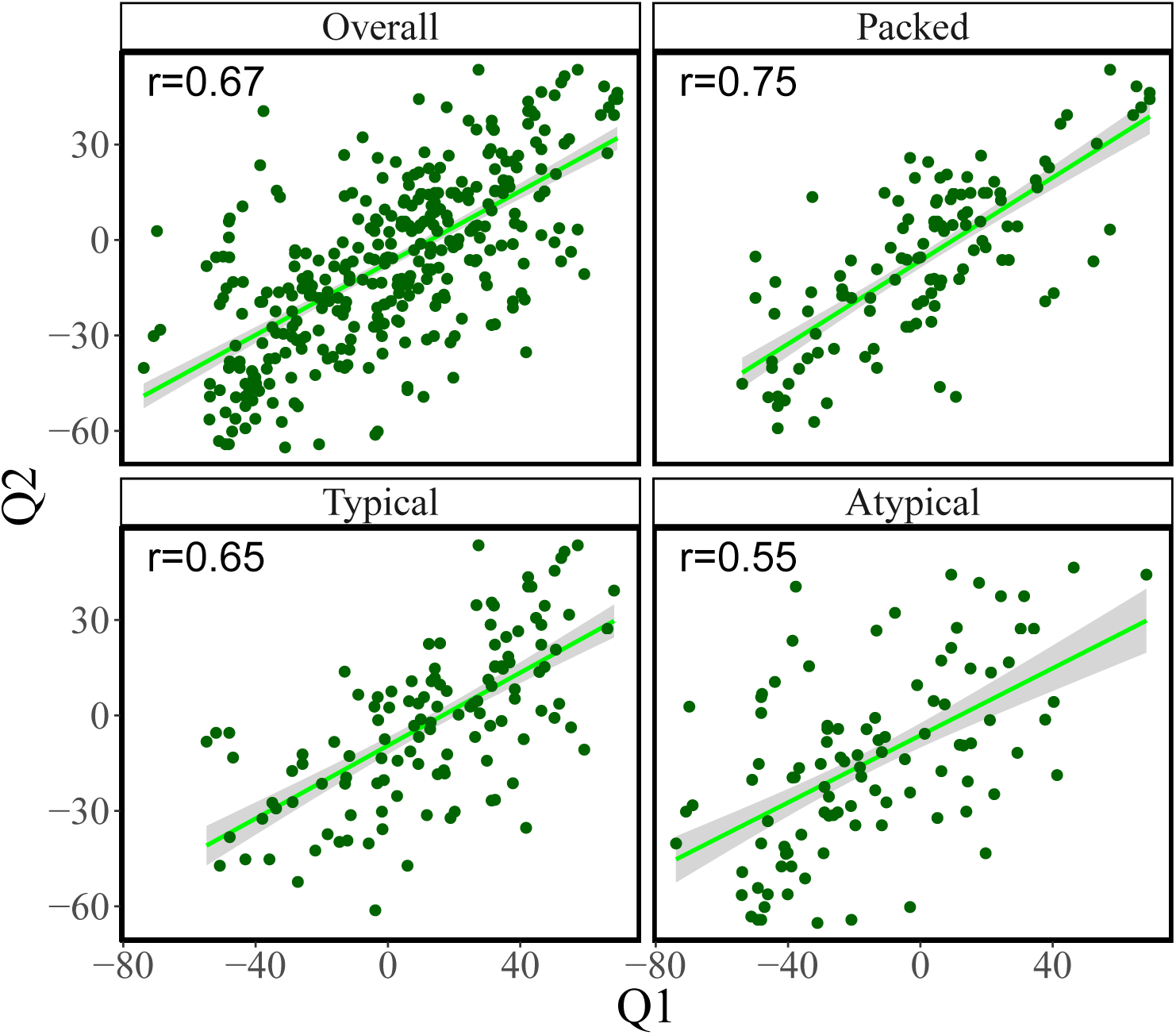
Trial-by-trial analyses of Experiment 1. Difference between *Q1* responses and true probability (as assessed by our MCMC model) plotted against the same quantity for *Q2*. Lines show the least-squares fit with standard error bands.

We found significant positive correlations between the two queries in all conditions when aggregating across participants (average correlation: *r* = 0.67, *p* < 0.01). The same conclusion can be drawn from analyzing correlations within participants and then testing the average correlation against 0 (*r* = 0.55, *p* < 0.01). Moreover, the within-participant effect size (the response difference between the unpacked conditions and the packed condition) for *Q1* was correlated with responses to *Q2* for both atypical (*r* = 0.35, *p* < 0.01) and typical (*r* = 0.21, *p* < 0.05) unpacking conditions. This means that participants who showed greater subadditivity or superadditivity for *Q1* also showed correspondingly greater effects for *Q2*.

### Discussion

Experiment 1 established a memory effect in probabilistic inference: answers to a query are influenced by answers to a previous query, thereby providing evidence for amortization. In particular, both a sub- and a superadditivity effect induced at *Q1* carried over to *Q2*, and participants showing stronger effects sizes for both sub- and superadditivity for *Q1* also showed greater effects for *Q2*.

## Experiment 2

Our next experiment sought to discriminate between sample-based and summary-based amortization strategies. We follow the logic of the simulations shown in Figure 2, manipulating cognitive load at *Q1* and *Q2* in order to exogenously control the number of samples (see Dasgupta, Schulz, & Gershman, 2017; Thaker et al., 2017, for a similar approach).

In addition to cognitive load, we manipulate the “overlap” of *Q1* with *Q2*, by creating a new set of “no overlap” queries with no overlap between the hypothesis spaces of the query pairs. We predicted that we would only see a memory effect for queries with overlapping pairs. This manipulation allows us to rule out an alternative trivial explanation of our results: numerical anchoring (high answers to the first query lead to high answers to the second query). If the apparent memory effect was just due to anchoring, we would expect to see the effect regardless of query overlap, contrary to our predictions.

### Participants

80 participants (53 males, mean age=32.96, SD=11.56) were recruited from Amazon Mechanical Turk and received $0.50 as a basic participation fee and an additional bonus of $0.10 for every on time response as well as $0.10 for every correctly classified digit during cognitive load trials.

### Procedure

The procedure in Experiment 2 was largely the same as in Experiment 1, with the following differences. To probe if the memory effects arise from reuse or from numerical anchoring, we added several **Q2** queries to the list shown in Table 1. These **Q2** queries have no overlap with the queried hypothesis for *Q1* (for example, ‘T or R’ instead of ‘C or R’ in the trial shown in the first row in Table 1). In other words, these queries could not be decomposed such that the biased samples from *Q1* be reflected in the answer to *Q2*, so the sub- and super-additive effects would not be seen to carry over to *Q2* were reuse to occur. We refer to these queries as “no overlap”, in contrast to the other “partial overlap” queries in which one of the letters overlapped with the previously queried letter. Half of the queries had no overlap and half had partial overlap, randomly interspersed. The stimuli used in Experiment 2 are shown in Table 2.

**Table 2.** Experimental stimuli and queries for Experiment 2.

| Cue | $Q1$ | $Q1$-Typical | $Q1$-Atypical | $Q2$-Partial overlap | $Q2$-No overlap |
| --- | --- | --- | --- | --- | --- |
| Table | C | chair, computer, curtain | cannon, cow, canoe | C or R | T or R |
| Telephone | D | display case, dresser, desk | drinking fountain, dryer, dome | D or L | G or L |
| Rug | B | book, bouquet, bed | bird, buffalo, bicycle | B or S | D or S |
| Chair | P | painting, plant, printer | porch, pie, platform | P or A | M or A |
| Sink | T | table, towel, toilet | trumpet, toll gate, trunk | T or E | F or E |
| Lamp | W | window, wardrobe, wine rack | wheelbarrow, water fountain, windmill | W or F | L or F |

To probe if the memory effect arises from reuse of generated samples (sample-based amortization) or reuse of summaries (summary-based amortization), we also manipulated cognitive load: on half of the trials, the cognitive load manipulation occurred at *Q1* and on half at *Q2*. A sequence of 3 different digits was presented prior to the query, where each of the digits remained on the screen for 1 second and then vanished. After their response to the query, participants were asked to make a same/different judgment about a probe sequence. Half of the probes were old and half were new.

We hypothesized that partial overlap would lead to a stronger amortization effects, whereas no overlap would lead to weaker effects. Furthermore, if participants are utilizing sample-based amortization, then cognitive load during *Q2* should increase the amortization effect: if more samples are generated during *Q1* (which are the samples that contain the sub- or superadditivity biases) and these samples are concatenated with fewer unbiased samples during *Q2*, then the combined samples will be dominated by biased samples from *Q1* and therefore show stronger effects. Vice versa, if participants are utilizing summary-based amortization, then cognitive load during *Q1* should increase the amortization effect: if less samples are generated during *Q1*, then a summary of those samples will inherit a stronger sub- or superadditivity effect such that the overall amortization effect will be stronger if the two summaries are combined (assuming that the summaries are combined with equal or close-to equal weights).

### Results

Analyzing only the queries with partial overlap (averaging across load conditions), we found that probability judgments for *Q1* were higher for unpacked-typical compared to packed conditions (a subadditivity effect; t(79) = 4.38, *p* < 0.001) and lower for unpacked-atypical compared to packed (a superadditivity effect; t(79) = —4.94, *p* < 0.001). These same effects occurred for *Q2* (unpacked-typical vs. packed: t(79) = 2.44, *p* < 0.01; unpacked-atypical vs. packed: t(79) = —1.93, *p* < 0.05).

We again calculated the difference between each participant’s response to every query during *Q1* and the overall mean response for the same query object in the packed condition. This difference was then used as the dependent variable in a linear mixed-effects regression model with participants and queried object as random effects and condition as fixed effect. The resulting estimates showed both a significant subadditivity (difference = 13.64 ± 1.57, t(396.95) = 8.70, *p* < 0.0001) and superadditivity (-14.90 ± 1.56, t(395.48) = -9.55, *p* < 0.0001) effect. Afterwards, we repeated the same analysis for responses to *Q2* (as in Experiment 1). This analysis again showed significant indicators of amortization as both the subadditivity (difference = 5.37 ± 1.34, t(398.01) = 4.02, *p* < 0.001) and the superadditivity effect (difference = -4.92 ± 1.336461, t(398.01) = -3.69, *p* < 0.001) were still present during *Q2*.

Next, we assessed how the memory effect was modulated by cognitive load and overlap (Figure 6). When cognitive load occurred during *Q2* and there was no overlap, none of the conditions produced an effect significantly different from 0 (all *p* > 0.5). When cognitive load occurred during *Q2* and there was partial overlap, only typically unpacked hypotheses produced an effect significantly greater than 0 (£(38) = 2.14, *p* < 0.05). When cognitive load occurred during *Q1* and there was no overlap, we found again no evidence for the conditions to differ from 0 (all *p* > 0.05). Crucially, if cognitive load occurred during *Q1* and there was partial overlap, both conditions showed the expected subadditive (£(38) = 4.18, *p* < 0.05) and superadditive (£(46) = —1.89, *p* < 0.05) effects. Moreover, calculating the average effect size of amortization for the different quadrants of Figure 6, the partial overlap-cognitive load at *Q1* condition produced the highest overall effect (d = 0.8), followed by the partial overlap-cognitive load at *Q2* condition (d = 0.56) and the no overlap-cognitive load at *Q1* condition (d = 0.42). The no overlap-cognitive load at *Q2* condition did not produce an effect greater than 0. Partial overlap trials were also more strongly correlated with responses during *Q1* than trials with no overlap (0.41 vs 0.15, £(157) = —2.28, *p* < 0.05).

**Figure 6.**
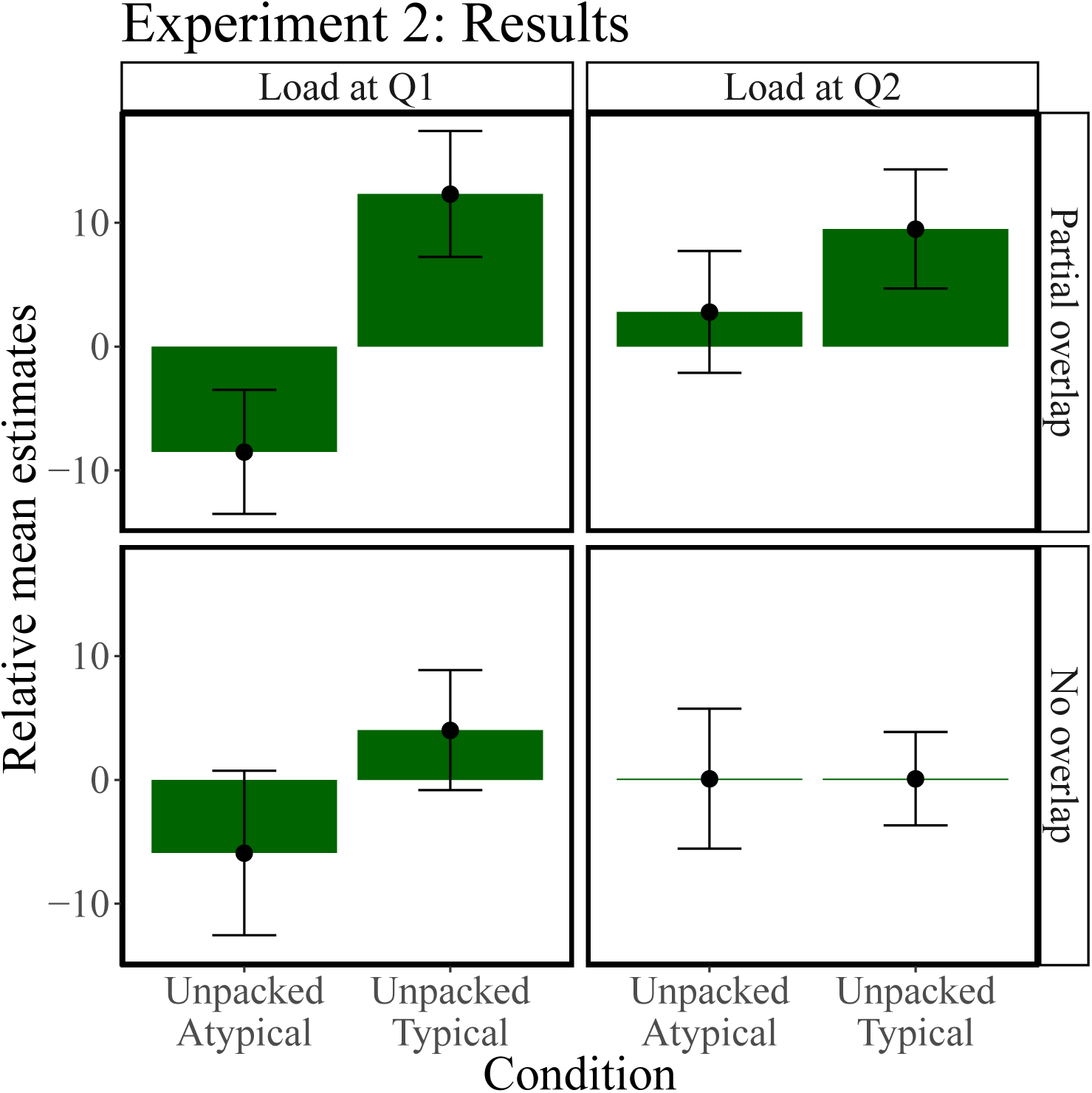
Experiment 2: Differences between *Q2* responses for each condition and an average packed baseline. A negative relative mean estimate indicates a superadditivity and a positive relative mean estimate a subadditivity effect. Error bars represent the standard error of the mean.

Next, we calculated the difference between all responses to *Q2* and the mean responses to *Q2* over queried objects provided that *Q1* was packed. This difference was enter into a linear mixed-effects regression that contained overlap, cognitive load, and the presentation format of *Q1* as fixed effects and participants and the queried objects as random effects. We then assessed the interaction between cognitive load and the sub- and superadditivity conditions while controlling for overlap. The resulting estimates showed that there was a significant subadditivity (difference = 5.25 ± 2.12, £(417.08) = 2.48 *p* < 0.05) but no superadditivity (difference = —3.19 ± 2.17, £(419.23) = —1.47, *p* = 0.17) effect when cognitive load was applied during *Q2*. Importantly, both the subadditivity (difference = 5.83 ± 2.25, £(418.91) = 2.59, *p* < 0.05) and the superadditivity (difference = —6.86 ± 2.21, £(419.80) = —3.102, *p* < 0.01) effect were present when cognitive load was applied during *Q1*. This finding points towards a larger amortization effect in the presence of cognitive load on *Q1*, thus supporting a summary-based over a sampled-based amortization scheme.

Further, on trials with cognitive load at *Q2*, participants were on average more likely to answer the probe correctly for partial overlap trials compared to no overlap trials (t(36) = 3.16, *p* < 0.05). This is another signature of amortization: participants are expected to have more resources to spare for the memory task at *Q2* if the computations they executed for *Q1* are reusable in answering *Q2*. This also indicates that these results cannot be explained by simply initializing the chain for *Q2* where the chain for *Q1* ended, which would not have affected the required computations.

Interestingly, there was no evidence for a significant difference between participants’ responses to *Q2* under cognitive load in Experiment 2 as compared to participants’ responses to *Q2* in Experiment 1 when no cognitive load during both *Q1* or *Q2* was applied in Experiment 1 (t(314) = —1.44, *p* = 0.15).

Finally, we assessed how much the difference between responses for *Q1* and the true underlying probabilities were predictive of the difference between responses for *Q2* and the true underlying probabilities (Figure 7). There was a strong correlation between responses to *Q1* and *Q2* over all conditions (r = 0.41, *p* < 0.001), for the packed (r = 0.44, *p* < 0.001), the typically unpacked (r = 0.36, *p* < 0.01), as well as the atypically unpacked condition (r = 0.40, *p* < 0.01). Moreover, the differences of *Q1* and *Q2* responses from the true answer were also highly correlated within participants (mean *r* = 0.31, *p* < 0.01) and participants who showed stronger subadditivity or superadditivity effects for *Q1* also showed stronger effects for *Q2* overall (r = 0.31, *p* < 0.001), for the superadditive (r = 0.3, *p* < 0.001), and for the subadditive condition (r = 0.29, *p* < 0.001). This replicates the effects of amortization found in Experiment 1.

**Figure 7.**
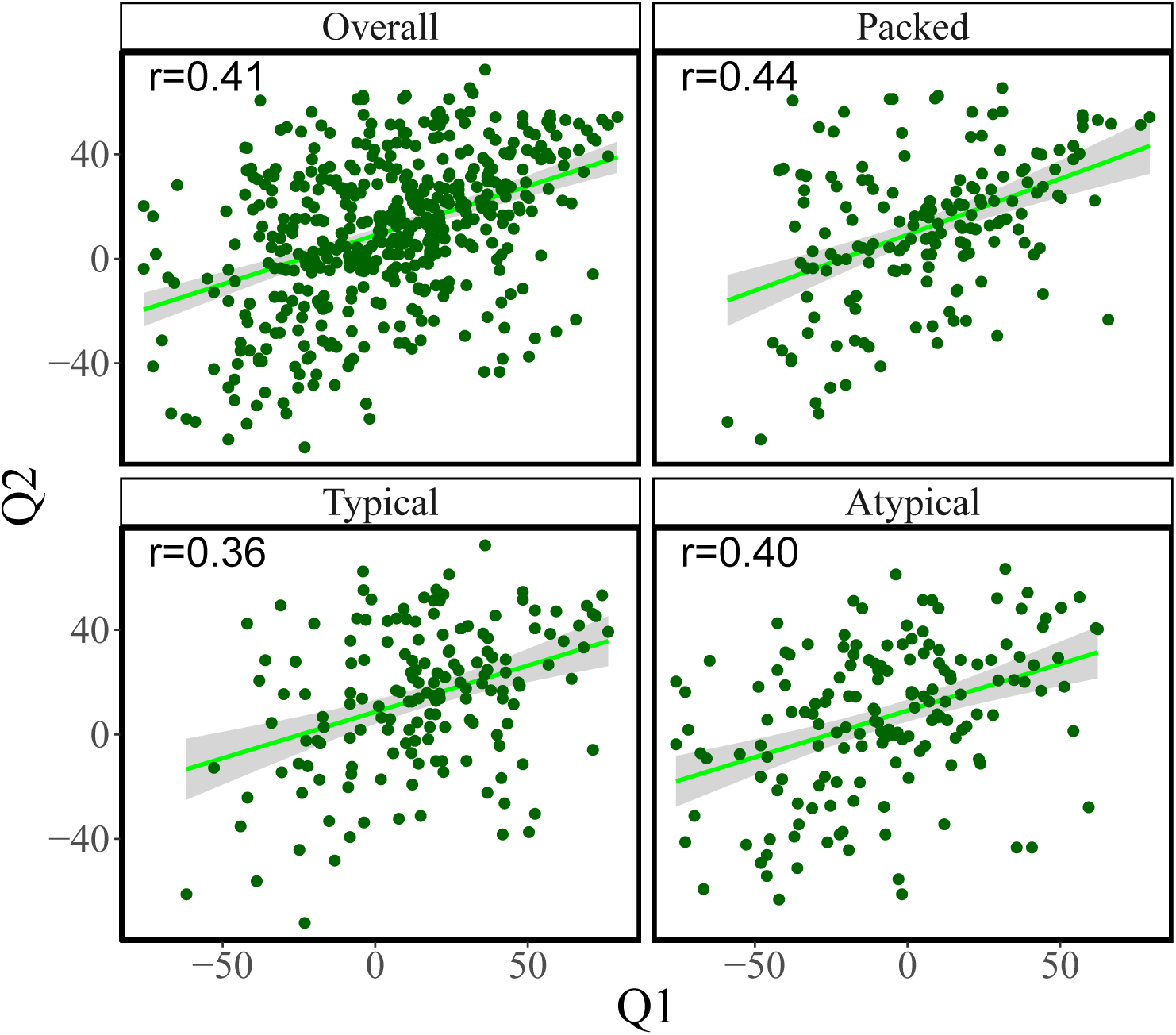
Trial-by-trial analyses of Experiment 2. Relationship between difference between *Q1* responses and true probability (as assessed by our MCMC model) and *Q2* responses and true probability. Lines show the least-squares fit with standard error bands.

### Discussion

Experiment 2 extended the findings of Experiment 1, suggesting constraints on the underlying amortization strategy. Participants exhibited an intricate pattern of sensitivity to cognitive load and query overlap. Based on our simulations (Figure 2), we argue that the effect of cognitive load at *Q1* on *Q2* responses is more consistent with summary-based amortization than with sample-based amortization. Summary-based amortization is less flexible than sample-based amortization, but trades this inference limitation for an increase in memory efficiency, and is thus consistent with the idea that humans adopt cost-efficient resource-rational inference strategies (Gershman et al., 2015; Griffiths, Lieder, & Goodman, 2015; Lieder et al., 2017a). Further supporting this idea is our finding that performance on the secondary task was better in the partial overlap conditions, indicating that more resources are available when computations can be amortized.

Our design allowed us to rule out a numerical anchoring effect, whereby participants would give high answers to the second query if they gave high answers to the first query. This effect should be invariant to the extent of overlap of the queried hypothesis spaces, but contrary to the anchoring hypothesis, the memory effect was stronger in the high overlap condition.

## Experiment 3

In this experiment we try to further probe the strategic nature of amortization. So far, all generated hypotheses have been reusable, since both queries probe the same probability distribution, conditioned on the same cue object. By changing the cue object between *Q1* and *Q2* and manipulating the similarity between the cues, we can control how reusable the computations are. Note that this is in contrast to the notion of “overlap” in Experiment 2 where all the samples from *Q1* are always “reusable” in *Q2* since both query the same probability distribution, but in the no overlap conditions, the queried hypotheses spaces do not overlap resulting in the biased samples from *Q1* not being reflected in *Q2* judgments. The notion of reusability now allows us to test whether or not reuse always occurs, or if it occurs preferentially when it is more applicable (i.e., under high similarity between cues).

### Participants

100 participants (41 females, mean age=35.74, SD=11.69) were recruited from Amazon Mechanical Turk and received $0.50 as a basic participation fee and an additional bonus of $0.10 for every on time response.

### Procedure

The procedure was similar to Experiments 1 and 2. The only difference was that participants were shown a new cue word for *Q2*, asking them to judge the probability of objects starting with the same letter as the letter from *Q1* with no conjunction of letters (i.e., same query space, full overlap). The query for *Q2* was always packed, as in previous experiments. The new cue words for *Q2* were generated to either have posterior with a low (similar cues) or a high (dissimilar cues) KL divergence from the *Q1* posterior. The range of KL divergences fell between 0 and 9; all similar cue words had conditional distributions with KL divergence of less than 0.1, and all dissimilar cue-words had a KL divergence of greater than 8.5. The exact KL divergences are reported in Table 3.

**Table 3.** Experimental stimuli and queries for Experiment 3. Kullback-Leibler (KL) divergence between the posteriors for Q1 and Q2 are shown in parentheses.

| **Cue1** | **Q1** | **Q1-Typical** | **Q1-Atypical** | **Cue2-sim (KL)** | **Cue2-diff (KL)** |
| --- | --- | --- | --- | --- | --- |
| Rug | B | book, bouquet, bed | bird, buffalo, bicycle | Curtain (0.080) | Car (8.658) |
| Chair | P | painting, plant, printer | porch, pie, platform | Book (0.031) | Road (8.508) |
| Sink | T | table, towel, toilet | trumpet, toll gate, trunk | Counter (0.001) | Sidewalk (8.503) |

### Results

Seven participants did not respond on time to more than a half of all queries and were therefore excluded from the following analysis.

We again found that probability judgments for *Q1* in the typically unpacked queries were higher than in the unpacked condition (subadditivity effect: t(92) = 4.67, *p* < 0.001) and that probability judgments in the atypically unpacked condition were lower than in the unpacked condition (superadditivity effect: t(92) = 3.25, *p* < 0.01).

Analyzing the probability judgments for *Q2*, we found a significant subadditivity effect ((t(92) = 2.28, *p* < 0.05) but not a significant superadditivity effect (56.06 vs. 55.31; t(92) = 0.07,p = 0.94).

As before, we calculated the difference between each participant’s response to every query during *Q1* and the overall mean response for the same query object in the packed condition. This difference was entered as the dependent variable into a linear mixed-effects regression model with participants and queried object as random effects and condition as fixed effect. The resulting estimates showed both a significant subadditivity (difference = 14.39 ± 1.97, t(189.84) = 7.31, *p* < 0.0001) and superadditivity (—13.72 ± 1.98, t(190.18) = —6.941, *p* < 0.0001) effect. Repeating this analysis for responses to *Q2* revealed a significant amortization effect for the the subadditivity (difference = 5.21 ± 1.90, t(191) = 2.74, *p* < 0.05) but not for the superadditivity condition (difference = —2.49 ± 1.91, t(191.52) = —1.303 *p* = 0.19).

**Figure 8.**
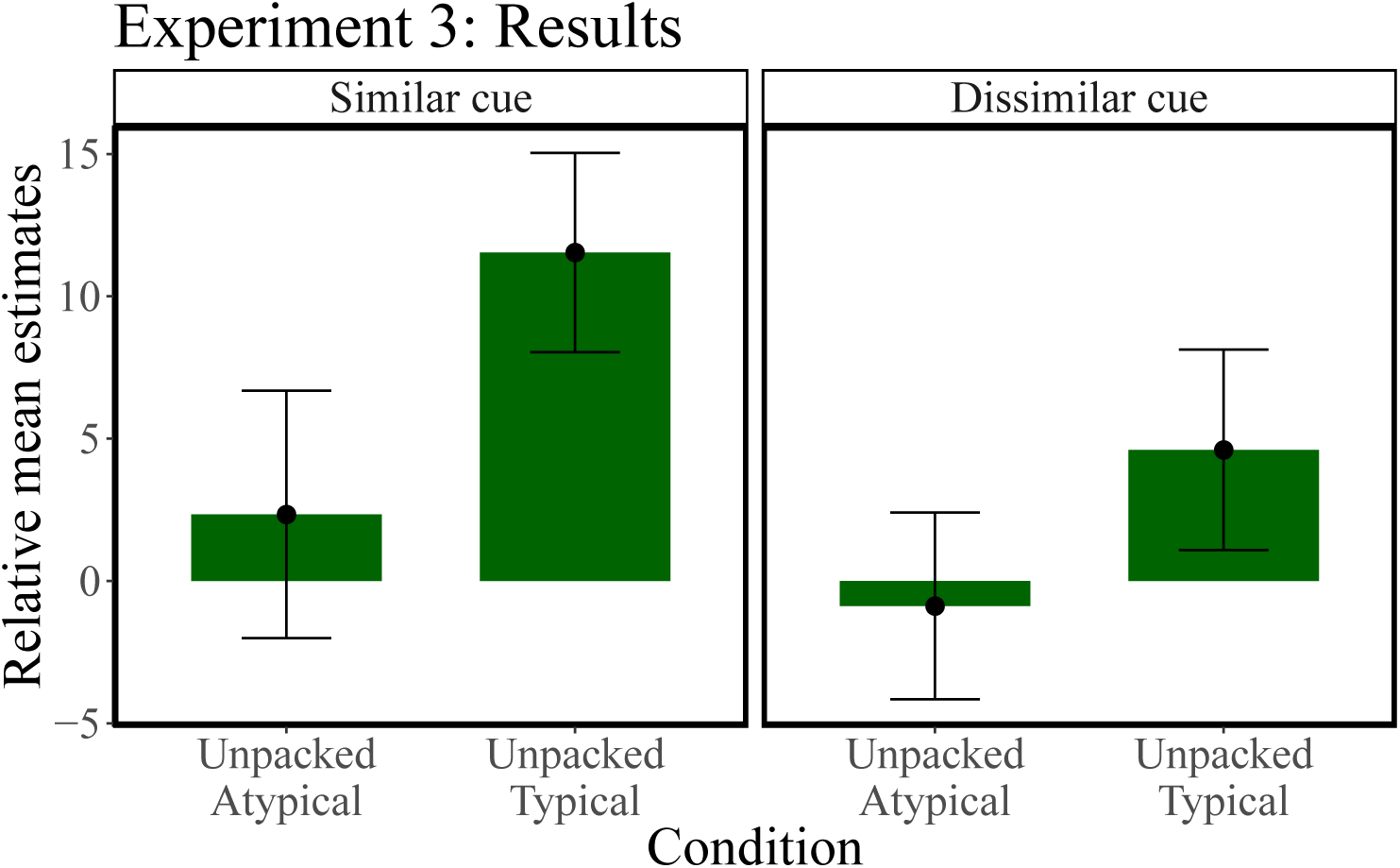
Experiment 3: Differences between *Q2* responses for each condition and an average packed baseline. A negative relative mean estimate indicates a superadditivity and a positive relative mean estimate a subadditivity effect. Error bars represent the standard error of the mean.

For the dissimilar cues, we did not find statistical evidence for an effect of subadditivity (£(49) = 1.31, *p* = 0.19) or superaditivity(£(47) = —0.27, *p* = 0.79). However, for the similar cues at *Q2*, the effect for the typically unpacked condition was significantly different from 0 (subadditivity effect: £(47) = 3.30, *p* < 0.01), whereas there was again no superadditivity effect (£(48) = 0.54, *p* = 0.59). The difference between the size of the subadditivity effect was marginally bigger for the similar cues as compared to the dissimilar cues (£(36) = 1.83, *p* = 0.06) and the overall effect size of the similar cues was *d* = 0.17, whereas the effect size for the dissimilar cues was *d* = 0.11.

The difference between judgments and the true probabilities was correlated between *Q1* and *Q2* (r = 0.34, *p* < 0.001), for the packed (r = 0.43, *p* < 0.001), the typically unpacked (r = 0.43, *p* < 0.001), but not the atypically unpacked condition (r = 0.20, *p* = 0.3); see Figure 9. Participants who showed higher subadditivity or superadditivity effects for *Q1* also showed higher effects for *Q2* overall (r = 0.29, *p* < 0.001), for the typically unpacked condition (r = 0.39, *p* < 0.001), but not for the atypically unpacked condition (r = 0.11, *p* = 0.29).

**Figure 9.**
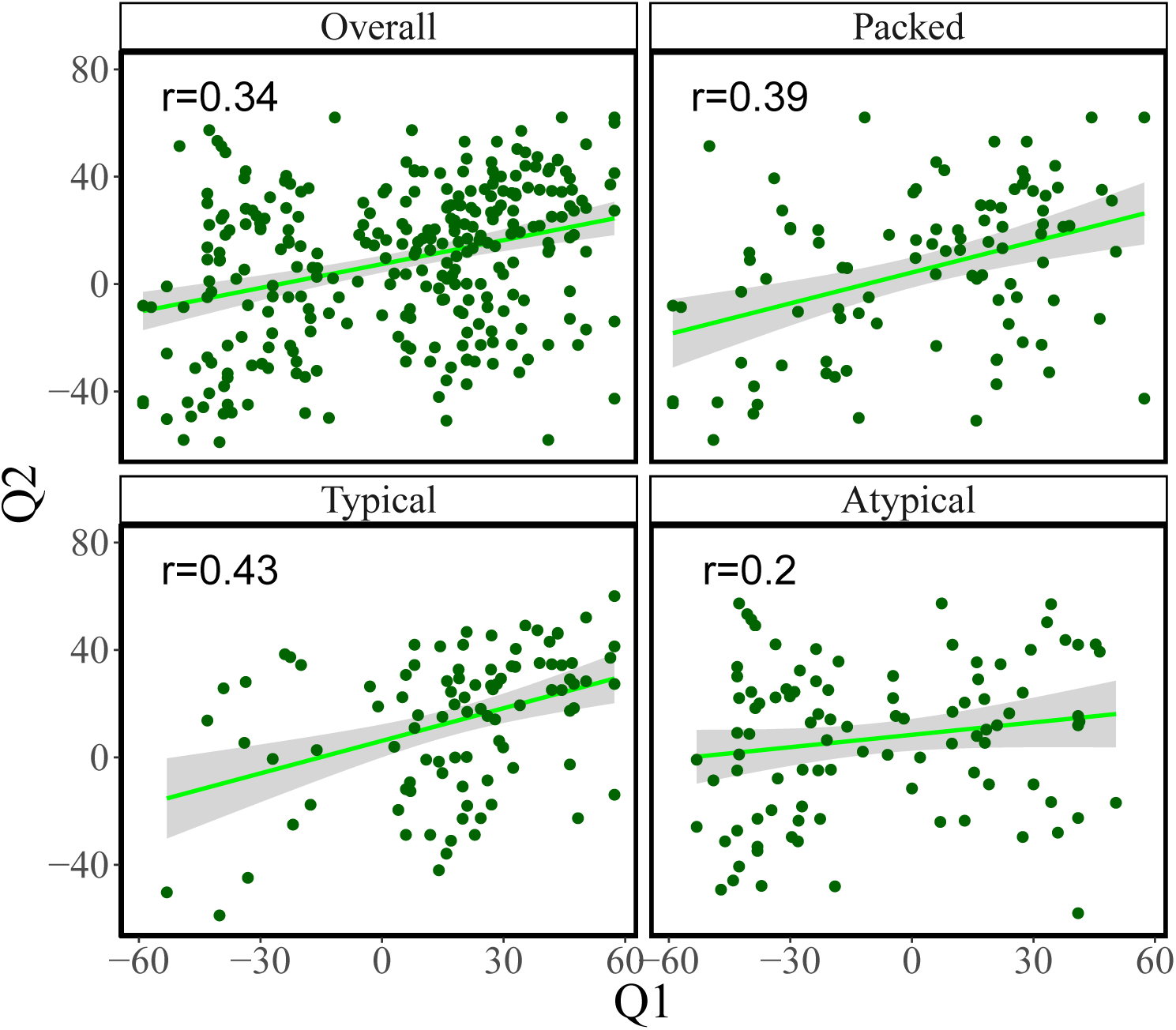
Trial-by-trial analyses of Experiment 3. Relationship between difference between *Q1* responses and true probability (as assessed by our MCMC model) and *Q2* responses and true probability. Lines show the least-squares fit with standard error bands.

### Discussion

Experiment 3 assessed the strategic nature of amortization by manipulating the similarity between cues, which presumably affected the degree to which amortization is useful. We found a stronger subadditivity effect for similar cues compared to dissimilar cues, indicating that reuse is at least partially sensitive to similarity.

An unexpected finding was that while the superadditivity effect in aytpically-unpacked *Q1* was significant, neither the memory-based superadditivity effect (in *Q2*) nor correlations across the queries for atypically-unpacked *Q1* were significant. This indicates that the answers to the atypically-unpacked *Q1* are not detectably being reused in *Q2* in this experiment. However, in Experiments 1 and 2, the atypically-unpacked answers seem to be reused (as indicated by a robust memory-based superadditivity effect, and correlations across the queries) *when the cue object remains the same*. This may be because the extent of rational reuse here (where the cues change) is smaller than in previous experiments (where the cues remained the same) and therefore harder to detect. Another possible explanation for this difference in Experiment 3 is that changing the cue object engages a sort of reset, and allows people to ignore low-probability/bad samples such as the ones occurring in the atypically-unpacked condition, and therefore dismiss the stored answers to atypically-unpacked *Q1*. Further research is needed to investigate this possibility.

## General Discussion

We tested a model of amortized hypothesis generation across 3 experiments and found that participants not only exhibited subadditive and superadditive probability judgments in the same paradigm (replicating Dasgupta, Schulz, & Gershman, 2017), but also carried over these effects to subsequent queries—a memory effect on inference. Experiment 2 demonstrated that this memory effect is some function of the hypotheses generated in the first query and made some inroads into trying to understand this function. We found that the effect is stronger when cognitive load is applied to the first query, suggesting that the memory effect is driven by a form of summary-based amortization, whereby a summary statistic of the first query is computed from the samples and then reused to answer subsequent queries, provided they can be expressed in terms of previous computations. Summary-based amortization gives up some flexibility (compared to reusing the raw samples generated by past inferences), in order to gain memory-efficiency. Experiment 3 demonstrated that the memory effect selectively occurs when the queries are similar, indicating that reuse is deployed specifically when it is likely to be useful.

Building on earlier results (Gershman & Goodman, 2014), our findings support the existence of a sophisticated inference engine that adaptively exploits past computations. While reuse can introduce error, this error may be a natural consequence of a resource-bounded system that optimally balances accuracy and efficiency (Gershman et al., 2015; Griffiths et al., 2015; Lieder et al., 2012; Vul et al., 2014). The incorporation of reuse into a Monte Carlo sampling framework allows the inference engine to preserve asymptotic exactness while improving efficiency in the finite-sample regime.

### Related work

This work fits into a larger nexus of ideas exploring the role of memory in inductive reasoning. Heit, Hayes and colleagues have carried out a number of studies that make this link explicit (Hawkins, Hayes, & Heit, 2016; Hayes, Fritz, & Heit, 2013; Hayes & Heit, 2013; Heit & Hayes, 2011). For example, Heit and Hayes (2011) developed a task in which participants studied a set of exemplars (large dogs that all possess “beta cells”) and then on a test set of exemplars (consisting of large and small dogs) made either property induction judgments (“does this dog have beta cells?”) or recognition memory judgments (“did this dog appear in the study phase?”). The key finding was that property induction and recognition memory judgments were strongly correlated across items, supporting the hypothesis that both judgments rely on a shared exemplar similarity computation: test exemplars are judged to be more familiar, and have the same latent properties, to the degree that they are similar to past exemplars. Heit and Hayes showed that both judgments could be captured by the same exemplar model, but with a broader generalization gradient for induction.

Another example of memory effects on inference is the observation that making a binary decision about a noisy stimulus (whether dots are moving to the left or right of a reference) influences a subsequent continuous judgment about motion direction (Jazayeri & Movshon, 2007). Stocker and colleagues (Luu & Stocker, 2016; Stocker & Simoncelli, 2008) refer to this as “conditioned perception”’ or “self-consistent inference” because it appears as though observers are conditioning on their decision as they make a choice. Fleming and Daw (2017) have pushed this idea further, arguing that observers condition on their own confidence about the decision. Self-consistent inferences may reflect rational conditioning on choice or confidence information when a memory trace of the stimulus is unavailable or unreliable.

An intriguing explanation of order effects has been reported by Wang and colleagues (Wang & Busemeyer, 2013; Wang, Solloway, Shiffrin, & Busemeyer, 2014). The key idea, derived from a quantum probability model of cognition (see also Trueblood & Busemeyer, 2011), is that answering a question will cause the corresponding mental state to linger and thus “superpose” with the mental state evoked by a second question. This superposition gives rise to a particular symmetry in the pattern of judgments when question order is manipulated, known as the *quantum question order equality* (see Wang & Busemeyer, 2013, for details). Our amortization framework does not intrinsically make this prediction, but nor does it necessarily exclude it. Rather, we prefer to think about superposition states as arising from computational principles governing a computation-flexibility trade-off. Roughly speaking, states superpose in our framework because the inference engine is reusing information from past queries.

Recently, Costello and Watts (2018) pointed out that the quantum question order equality could arise from rational probabilistic reasoning corrupted by correlated noise. In particular, answers to a probabilistic query will be corrupted by samples retrieved recently to answer another probabilistic query (similar to the concept of “overgeneralization” in probabilistic estimation, as developed in Marchiori, Di Guida, & Erev, 2015). Costello and Watts (2018) view this as a kind of priming effect. Alternatively, correlated noise would arise in the amortized inference framework due to stochastic reuse. Thus, amortization might provide a complementary rational analysis for the “probability theory plus noise” model proposed by Costello and Watts (2018).

Most closely related to the present paper is the work of Dougherty and colleagues (Dougherty, Gettys, & Ogden, 1999; Dougherty & Hunter, 2003b, 2003a; Thomas, Dougherty, Sprenger, & Harbison, 2008; Thomas, Dougherty, & Buttaccio, 2014), who have pursued the idea that probability judgments depend on the generation of hypotheses from memory. In particular, they argue that subadditivity arises from the failure to generate hypotheses, much like the account offered by Dasgupta, Schulz, and Gershman (2017), and that this failure is exacerbated by cognitive load or low working memory capacity. The key difference from our account is the particular way in which memories are used to generate hypotheses. For combinatorial hypothesis spaces like the scene inference task used here and by Dasgupta, Schulz, and Gershman (2017), one cannot assume that all the relevant hypotheses are already stored in memory; rather, these must be generated on the fly—a function we ascribe to MCMC sampling. The present paper asserts a more direct role for memory within a sampling framework, by controlling the trade-off between computation and flexibility.

This trade-off mirrors a similar tension in reinforcement learning, where the goal is to estimate long-term reward (Daw, Gershman, Seymour, Dayan, & Dolan, 2011; Daw, Niv, & Dayan, 2005; Kool, Gershman, & Cushman, 2017). “Model-based” algorithms estimate long-term reward by applying tree search or dynamic programming to a probabilistic model of the environment. This is flexible, but computationally expensive. “Model-free” algorithms avoid this cost by directly estimated long-term rewards by interacting with the environment, storing these estimates in a look-up table or function approximator. This is computationally cheap but inflexible. In other words, model-free algorithms trade time for space, much in the same way that amortized inference uses memory to reduce the cost of approximate inference. Analogous to our proposed summary-based amortization strategy, recent work has suggested that model-free value estimates can be incorporated into model-based tree search algorithms (Keramati, Smittenaar, Dolan, & Dayan, 2016), thus occupying a middle ground in the time-space trade-off.

### Future directions

Our work has focused on fairly simple forms of amortization. There exists a much larger space of more sophisticated amortization strategies developed in the machine learning literature (e.g., Rezende et al., 2014; Stuhlmüller et al., 2013) that we have not yet explored. Finding behaviorally distinguishable versions of these algorithms is an interesting challenge. These versions could take the form of reuse in much more abstract ways, such as developing strategies and heuristics, instead of just local reuse in a sequence of queries. We believe that further examining established effects of heuristics and biases through the lens of computational rationality will continue to produce interesting insights into principles of cognition.

More broadly, we are still lacking a comprehensive, mechanistic theory of amortized inference. What objective function is being optimized by amortization? How are the computational trade-offs managed algorithmically? What are the contributions of different memory mechanisms (episodic, semantic, procedural, etc.)? Answering these questions will require a more general theoretical treatment than the one offered here. Nonetheless, our experiments provide important constraints on any such theory.

### Acknowledgments

We thank Kevin Smith for helpful comments and discussions. This material is based upon work supported by the Center for Brains, Minds and Machines (CBMM), funded by NSF STC award CCF-1231216. E.S. was supported by a postdoctoral fellowship from the Harvard Data Science Initiative.

## Appendix Two Re-use Schemes

The two schemes for re-use described in Figure 2, *summary-based* and *sample-based* amortization, are described below in greater detail.

In *sample-based amortization*, we simply add samples generated in response to one query (*Q1*) to the sample set for another query (*Q2*). So if *N_1_* samples were generated in response to *Q1*, and *N_2_* new samples are generated in response to *Q2*, in the absence of amortization, the responses to the two queries *Q1* and *Q2* would be generated as follows:

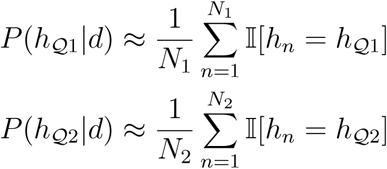

Under the sample-based amortization scheme, the response to *Q2* is given by a calculation carried out over all N + N2 equally weighted samples.

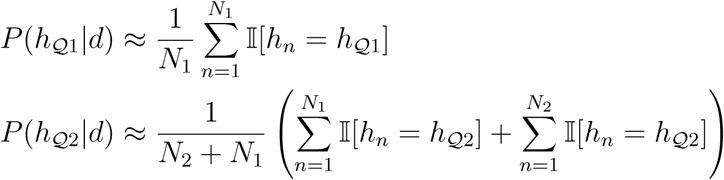

Under this scheme, all the computations carried out for *Q1* are available for flexible reuse in the computation for *Q2*.

In *summary-based amortization,* we reuse a summary statistic computed from *Q1*. This strategy is only applicable to problems where the answer to *Q2* can be expressed as the composition of the answer to *Q1*, and an additional simpler computation. For example if *Q1* is “What is the probability that there is an object starting with a C in the scene?”, *Q2* could be “What is the probability that there is an object starting with a C or an R in the scene?”. In this case, the N_1_ samples generated in response to *Q1* are summarized into one probability (“the probability of an object starting with C”), *N2* new samples are generated in response to a simpler query (“the probability of an object starting with R”), and these two numbers are then composed (in this case simply added) to give the final estimate for *Q2* (“the probability of an object starting with C or R”).

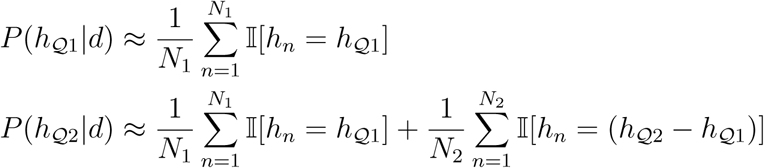

Under this scheme, only the final product of the computation carried out for *Q1* is reused in the calculations for *Q2*.

## Footnotes

1 More formally, this is known as variational inference (Jordan, Ghahramani, Jaakkola, & Saul, 1999), where the divergence is typically the Kullback-Leibler divergence between the approximate and true posterior. Although this divergence cannot be minimized directly (since it requires knowledge of the true posterior), an upper bound can be tractably optimized for some classes of approximations.

2 Our findings do not strongly depend on the use of the KL divergence measure and all of our qualitative effects remained unchanged when we applied a symmetric distance measure such as the Jensen-Shannon divergence.

